# MHChron: diversity-balanced dataset design for robust peptide–MHC binding prediction across MHC class I and II

**DOI:** 10.64898/2026.07.30.741752

**Authors:** Marta Chronowska, Eugene Shrimpton-Phoenix, Tadas Kluonis

## Abstract

Accurate prediction of peptide–MHC (pMHC) binding is central to immunogenicity assessment, yet many existing predictors are trained and evaluated on narrow allele sets and restricted peptide-lengths. Here, we present MHChron, a unified pMHC binding prediction framework predicated on systematic data curation, meticulous engineering of dataset balance and diversity, and rigorous evaluation through careful splits controlling for data leakage. We assemble one of the most diverse pMHC training dataset reported to date, integrating publicly available binding data across a broad allele coverage (class I n=214, class II n=98) and peptide length range (from 8 to 36 residues). Using a focused and carefully sampled subset of this dataset, we train complementary sequence-based and structure-aware models and test them under increasingly stringent generalisation regimes. Both models achieve consistently strong performance, outperforming the evaluated state-of-the-art predictors despite being trained on numerically fewer data points. Notably, the structure-aware model did not consistently surpass the sequence-based model, except under the most demanding setting of extrapolation to unseen allele clusters, suggesting that performance gains stem primarily from dataset diversity and rigorous evaluation rather than architectural complexity. Sequence-based MHChron is released with reproducible installation and an automated whole-protein screening pipeline, enabling broad and practical use.

## 1 Introduction

### 1.1 Biological basis of immunogenicity

The immune system is the body’s primary defence mechanism, continuously distinguishing between harmless self and potentially harmful non-self molecules [1]. When this decision fails, the consequences range from autoimmune disease to life-threatening drug reactions [2–6]. Predicting these failures computationally is therefore of crucial importance to allow safe development of therapeutics, personalised immunotherapy, gene therapy, organ transplantation, and efficient pandemic response.

The immune defence is mediated by two complementary branches: a rapid, non-specific innate system, and the slower, but more targeted adaptive system [7]. The cells of the adaptive immune system include B cells and T cells, which differ in how they detect molecular targets. B cells produce antibodies that bind intact three-dimensional biomolecular surfaces, such as exposed regions of proteins. In contrast, T cells do not recognise whole proteins, but instead survey short peptide fragments presented on the surfaces of other cells [8].

The production and presentation of these peptides occurs through two principal pathways. Intracellular proteins are primarily processed through the major histocompatibility complex (MHC) class I pathway, in which proteins are degraded in the cytosol by intracellular proteolytic enzymes into peptide fragments, transported into the endoplasmic reticulum, and loaded onto class I MHC molecules (MHC-I) for presentation at the cell surface to cytotoxic T cells (CD8^+^). In parallel, proteins acquired predominantly from the extracellular environment are internalised by specialised antigen-presenting cells, enzymatically cleaved into peptides within endosomal compartments, and presented via MHC class II molecules (MHC-II) to helper T cells (CD4^+^). Although these routes differ in processing and transport mechanisms, both converge on a common requirement: only peptides capable of forming sufficiently stable peptide–MHC (pMHC) complexes reach the cell surface for T-cell surveillance [9]. Importantly, MHC binding does more than simply permit cell-surface presentation: it also directly shapes the molecular interface recognised by the T-cell receptor (TCR) by constraining the bound peptide into a specific conformation [10]. Productive T-cell activation therefore requires a precise three-way compatibility between the peptide, the MHC molecule, and the TCR (Fig. 1A).

The formation of immunologically relevant pMHC complexes represents only one step within a broader cascade that includes post-translational modifications, antigen processing, subcellular localisation and transport, pMHC complex stability, and TCR engagement [11–17]. Nevertheless, among the many processes governing immunogenicity, pMHC binding exerts the strongest selective constraint on which peptide fragments ultimately become visible to T cells. Consequently, prediction of pMHC binding likelihood from paired peptide and MHC sequences has become a central task in computational immunogenicity assessment.

### 1.2 Sources of complexity in peptide–MHC binding prediction

In humans, MHC proteins are termed *human leukocyte antigens* (HLAs) and are encoded within one of the most polymorphic regions of the genome. The diversity is organised across six classical transplantation loci: three for class I (HLA-A, -B, and -C) and three for class II (HLA-DP, -DQ, and -DR) [18]. Each of these loci comprises thousands of documented allelic variants, grouped into hundreds of families [19]. This extensive allelic diversity enables recognition of a broad repertoire of antigenic peptides across individuals, but creates a major computational challenge because each HLA allele exhibits a unique peptide-binding specificity that must be experimentally measured or learned from data.

Despite differences in antigen source and processing context, both MHC classes share an overall similar structural fold, with the peptide-binding groove composed of a curved *β*-sheet platform at the base, flanked by two *α* helices. The critical difference is that the MHC-I’s peptide-binding platform is composed of a single polymorphic heavy chain, whereas MHC-II employs two asymmetric chains (Fig. 1B) [20]. This architectural divergence introduces distinct sequence and structural determinants that complicate the development of unified computational representations across both MHC classes.

**Fig. 1.**
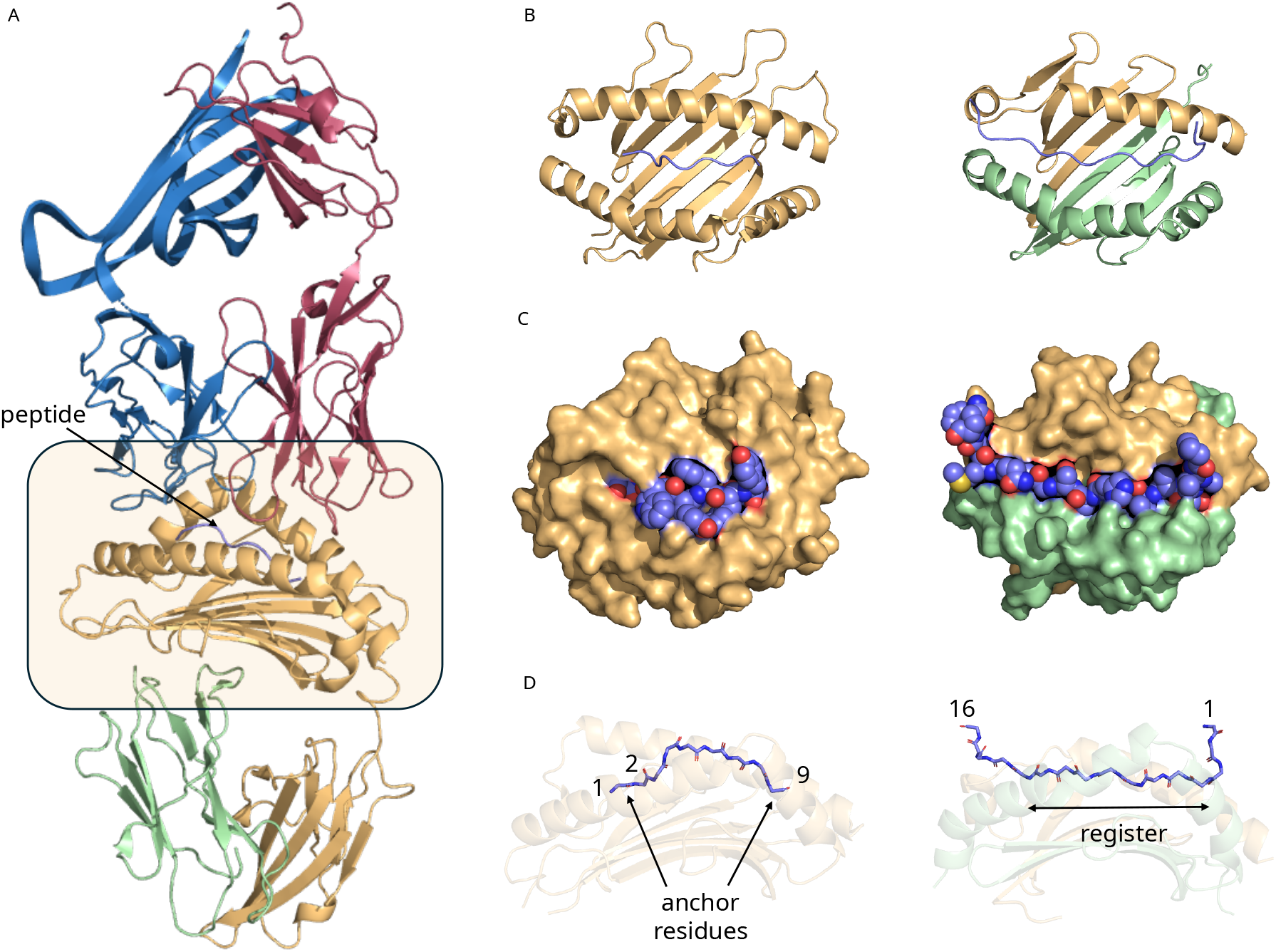
Structural basis of peptide presentation by MHC-I and MHC-II molecules and sources of binding-mode variability relevant to pMHC prediction. A) Structure of a TCR–peptide–MHC-I complex (PDB: 1G6R). The peptide-binding platform of the MHC molecule (yellow) constrains peptide conformation and contributes directly to the surface recognised by the T-cell receptor (TCR). TCR *α* chain = blue, TCR *β* chain = red; peptide = purple; MHC-I *α* chain = yellow; *β*2-microglobulin = green. B) Top-down views of the peptide-binding domains of MHC-I (left, PDB: 7LGD) and MHC-II (right, PDB: 7T6I). Despite differences in chain composition, both classes adopt a conserved architecture comprising a curved *β*-sheet floor flanked by two *α*-helices that form the peptide-binding groove. MHC-I consists of a single polymorphic *α* chain (yellow), whereas MHC-II is formed by two asymmetric chains (*α* in yellow, *β* in green). C) Surface representations of the same complexes shown in B), with peptides depicted as spheres. The MHC-I groove is closed at both termini, favouring shorter peptides, whereas the MHC-II groove is open-ended, allowing longer peptides to extend beyond the binding site. D) Side views with partially transparent MHC molecules highlighting peptide binding modes. MHC-I peptides (commonly 9 residues long) bind with conserved anchoring residues (e.g. P1, P2, and P9), often resulting in a bulged central conformation. In MHC-II, peptides bind via a typically nine-residue core (“register”) that lies flat within the groove. The example shown illustrates a non-canonical reverse-binding orientation, highlighting how multiple possible binding registers, register shifts, and occasional reverse-binding modes contribute to the complexity of pMHC binding prediction.

The two MHC classes differ further in the length and binding mode of the peptides they accommodate. Specifically, MHC-I molecules tend to bind shorter peptides (8-10 amino acids long) with termini buried in the closed binding groove, while MHC-II molecules typically associate with much longer peptides (13-25 amino acids long), due to the open conformation allowing flanking residues to protrude outwards (Fig. 1C) [20, 21]. Consequently, models must account for substantially different peptide-length distributions and binding geometries across the two MHC classes, limiting transferability of approaches developed for either class alone.

Both classes show plasticity in antigen presentation, with known examples of atypical peptide lengths, non-canonical binding modes (N- to C-terminus orientation relative to the *β*-sheet platform), or shifted registers (anchoring residues interacting with key pockets in the binding groove) (Fig. 1D) [21]. Together, these observations highlight that peptide–MHC binding cannot always be described by fixed positional rules, requiring prediction methods capable of capturing alternative binding conformations and context-dependent interactions.

### 1.3 Advances in computational prediction of peptide–MHC binding

Given the broad research and commercial importance of pMHC binding prediction, the field has witnessed sustained methodological innovation to address the extreme polymorphism of MHC molecules, variable peptide lengths, and sparse data coverage. In parallel, recent advances in protein representation learning - particularly protein language models (PLMs) such as ESM [22] and structure prediction tools such as AlphaFold2 (AF2) [23] - have expanded the toolkit for modelling protein-peptide interactions. While many established pMHC predictors do not directly incorporate these models, they have reframed how sequence and structural information can be encoded and integrated, prompting renewed comparison of sequence-based versus structure-aware approaches.

#### 1.3.1 Sequence-based peptide–MHC binding predictors

Sequence-based methods represent the earliest and most widely adopted type of pMHC binding predictors. These approaches typically represent peptides and MHC proteins using evolutionary features (through, for example, BLOSUM-derived matrices [17, 24–27]), categorical (“one-hot”) encodings [28–31], physicochemical descriptors [32, 33], or, more recently, PLM embeddings [34–38]. To meet the fixed-length input requirement of many machine learning models, these methods use a range of strategies for variable-length peptides, including length restriction [31, 38], sliding-window extraction [39, 40], padding [26, 34, 38, 41], alignment with insertions/deletions [25, 42], or other transformations [27]. MHC representations range from full-length sequences [14, 25, 29, 36, 39] to pseudosequences of putative contact residues [29, 38, 43]. The latter may be defined globally for an MHC class or selected for individual alleles using sequence- or structure-based analyses. Peptide and MHC encodings are either directly concatenated or separated by dedicated tokens inspired by natural language processing [34, 35, 37, 38].

Although many of these models report high AUROC values (often 94%–99%) at publication [27, 28, 44], subsequent studies revealed lower performance [45], discordant predictions across tools [46], and limited generalisability to unseen peptide lengths or under-represented alleles [47]. These findings suggest that, while effective at capturing statistical patterns from large immunopeptidomics datasets, sequence-centric models remain sensitive to imbalances in allele coverage and peptide-length distributions.

#### Structure-based peptide–MHC binding predictors

In contrast to the high sequence polymorphism of HLA alleles, the three-dimensional architecture of the MHC peptide-binding groove is highly conserved [48, 49]. Structure-based approaches therefore provide several conceptual advantages: they capture physicochemical interaction patterns and interface geometry in a more class-agnostic manner, offer improved interpretability through explicit atomic contacts and energy terms, and potentially increase data efficiency by leveraging shared binding modes across alleles.

Early physics-based structure-prediction pipelines relied on template superposition, docking, conformational sampling, and physics-derived scoring functions [50, 51]. These tools demonstrated the feasibility of the approach, but were limited by template availability, rigid-backbone approximations, and poor coverage of peptide lengths and MHC alleles. Subsequent methods extended this foundation by incorporating refined template selection with explicit anchor-restrained loop modelling [48, 52–55]. Despite gains, such pipelines remained computationally intensive, with accuracy declining markedly for longer peptides. More recent strategies have adapted AI-based structure prediction methods through specialised fine-tuning, enabling faster modelling across broader ranges of MHC sequences and peptide lengths [56–58]. More lightweight and efficient neural models continue to emerge, although these often reintroduce restrictions on peptide length or allele coverage [59–61], highlighting the continued evolution of the field and the recurring trade-off between accuracy, efficiency and applicability.

Despite these advances, several challenges remain, including limited experimental structural coverage, high computational costs, register identification errors (particularly for MHC-II), and difficulties in capturing conformational dynamics relevant to T cell recognition. Together, these factors constrain both the accuracy of predicted pMHC structures and the biological insights that can be derived from them [58, 62, 63].

### 1.4 Biases in peptide-MHC binding data

Large-scale datasets enabled by extensive data curation efforts from repositories such as the Immune Epitope Database and Analysis Resource (IEDB) [64] and IPD-IMGT/HLA Database [65] have substantially advanced our understanding of pMHC interactions. However, like all biological datasets, these resources inevitably reflect biases introduced by technical limitations and study design.

Despite nearly 43,000 HLA allelic sequences having been identified to date, peptide-binding data exist for fewer than 1% of them. Moreover, recent analyses have highlighted “alarming disparities” in HLA-associated data availability across different racial and ethnic populations, suggesting that current datasets may inadequately represent global HLA diversity and may reinforce existing inequalities in immunological research [66].

Beyond demographic representation, research priorities also shape the composition of available datasets. Experimental studies frequently focus on a limited number of common HLA alleles, particularly those with established relevance to therapeutic applications. Similarly, many generate peptide libraries with limited sequence diversity [67, 68]. These choices are scientifically understandable given experimental cost and feasibility, but may limit the ability of computational models to generalise beyond well-characterised regions of peptide and HLA sequence space.

Technical constraints further contribute to dataset bias. For example, mass spectrometry-based immunopeptidomics does not provide a direct measurement of peptide-binding affinity alone; instead, observed peptide presentation reflects a complex combination of antigen processing factors and experimental detectability. Physicochemical properties such as peptide hydrophobicity, ionisation efficiency, and pMHC complex stability can influence which ligands are experimentally recovered and quantified [69–71].

Consequently, training examples remain unevenly distributed across peptide lengths and HLA allele groups, and many models may preferentially exploit dataset-specific statistical regularities that correlate with binding measurements but fail to capture transferable biochemical constraints [72]. We therefore hypothesise that rigorous dataset curation, transparent evaluation across diverse HLA and peptide contexts, and biologically informed representations capturing structural and physicochemical constraints of pMHC interactions will be more critical for achieving robust generalisation than increasing model complexity alone.

### 1.5 Present study

In this work, we present MHChron, a pMHC binding prediction framework built on a balanced and highly diverse dataset spanning both MHC-I and MHC-II molecules. The dataset covers the largest number of unique alleles and the broadest peptide-length range reported for training pMHC binding predictors to date, with uniform representation across peptide lengths, alleles, and binding labels, to promote generalisation and enable realistic benchmarking.

Using this dataset, we train and compare two complementary models of matched expressive capacity: a structure-aware GATv2Conv [73] graph neural network (GNN) and a graph-free, sequence-only SetTransformer [74]. Both models were rigorously evaluated, including peptide- and allele-cluster-held-out splits, to explicitly probe generalisation beyond memorisation; we report AUROC, AUPRC and F1, along with locus- and peptide-length binned statistics. Across these settings, both models achieve strong and stable performance, suggesting that careful dataset design and training strategies yield greater gains than added model complexity alone.

The key contributions of this work are: methods for curating compact, high-quality training datasets; (2) splitting strategies that preserve balance across train, validation, and test partitions; (3) practical framework for integrating sequence and structural information; and (4) design principles for universal pMHC binding predictors. We release the sequence-based predictor as free user-facing software with streamlined installation, automated screening workflows, and interpretable visual outputs at https://github.com/wells-wood-research/MHChron (Fig. SI1 and SI2).

## 2 Results

### 2.1 Dataset preparation

The final dataset used for model development was balanced with respect to sequence composition, peptide length, allele representation, and binding labels, while maintaining a constrained total sample size.

The raw dataset comprised 18,768,920 peptide–MHC pairs, including 4,692,230 experimentally determined measurements and 14,076,690 decoy pairs. These spanned 1,064,430 unique peptide sequences and 315 MHC alleles (217 MHC-I *α*-chains, 19 MHC-II *α*-chains, and 79 MHC-II *β*-chains). The raw data contained approximately equal numbers of MHC-I and MHC-II examples. Peptide-length distributions were highly skewed, with 9-mers dominating the MHC-I data and 15-mers overrepresented in the MHC-II data.

The raw dataset was first subsampled to 50,000 entries, after which entries containing unnatural amino acids were removed. Structures were then generated using TFold, an AF2-based pipeline [58], yielding a final dataset of 42,012 pMHC complexes. This dataset was used for all subsequent analyses. It comprised 32,805 unique peptide sequences and 296 unique MHC peptide-binding domain sequences following TFold truncation of the full-length allele sequences (see Methods section 5.1). These corresponded to 214 MHC-I *α*-chain alleles, 19 MHC-II *α*-chain alleles, and 79 MHC-II *β*-chain alleles. Peptide lengths were uniformly distributed from 8 to 36 residues, in contrast to the strong enrichment of 9-mers and 15-mers in the source data. Binders and non-binders were represented in equal proportions across peptide lengths and MHC classes. MHC-II accounted for 75% of data points in the final dataset.

Changes in dataset composition throughout the curation pipeline are summarised in Fig. 2 and Fig. SI3.

**Fig. 2.**
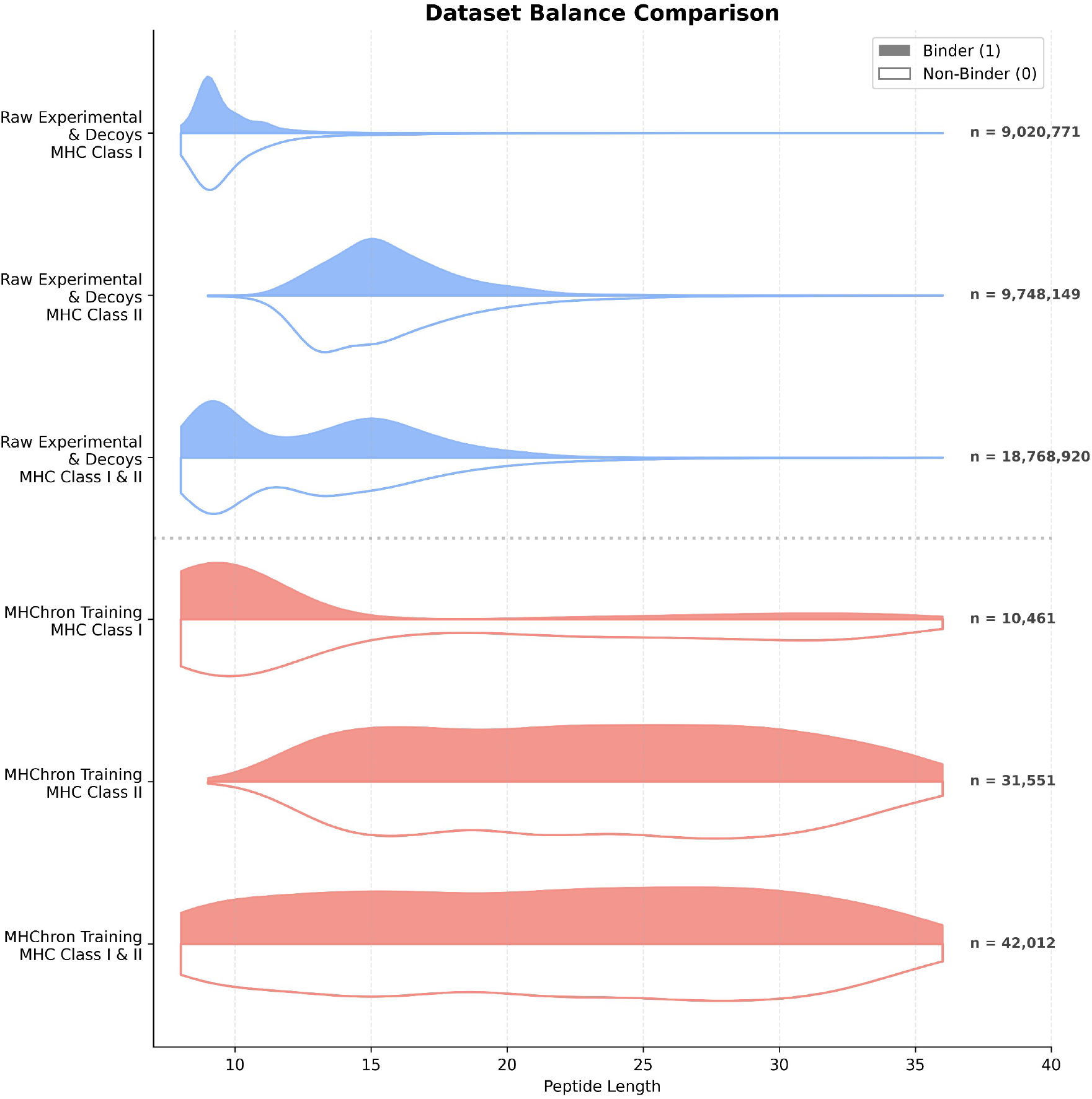
Comparison of peptide-length distributions between raw and MHChron training datasets. The raw experimental and decoy data (top, blue) exhibits a significant skew towards canonical lengths (i.e. 9-mers for MHC-I and 15-mers for MHC-II). In contrast, the MHChron training dataset (bottom, red) employs a strategic subsampling approach to achieve a balanced distribution across peptide lengths (8–36), binding labels (binder vs. non-binder), and MHC classes, mitigating length-based inductive bias during model training.

### 2.3 Model development

To guide model design, we trained multiple GATv2Conv-based GNNs, each using a different controlled combination of node and edge features to assess their relative contributions to predictive performance. These exploratory experiments were performed using the exploratory split described in Section SI4. To disentangle the contributions of pretrained PLM embeddings (ESM) from those of the explicit graph structure, we implemented a graph-free SetTransformer with representational capacity matched to the GATv2Conv architecture. The architectures of both models are shown in Fig. 3. Across the tested configurations, models incorporating per-residue ESM embeddings consistently achieved the strongest predictive performance, while additional or alternative node and edge representations provided little benefit. Balancing predictive performance against model size, computational complexity and feature-generation cost, we selected the GNN using ESM node embeddings and distance-based edges as the primary structure-aware model for all subsequent analyses (hereafter MHChron-*edgy* ), alongside the ESM-based sequence-only SetTransformer as a lightweight graph-free alternative (hereafter MHChron-*light* ). Full details of this exploratory analysis are provided in the Supplementary Information (Section SI3, Fig. SI4).

**Fig. 3.**
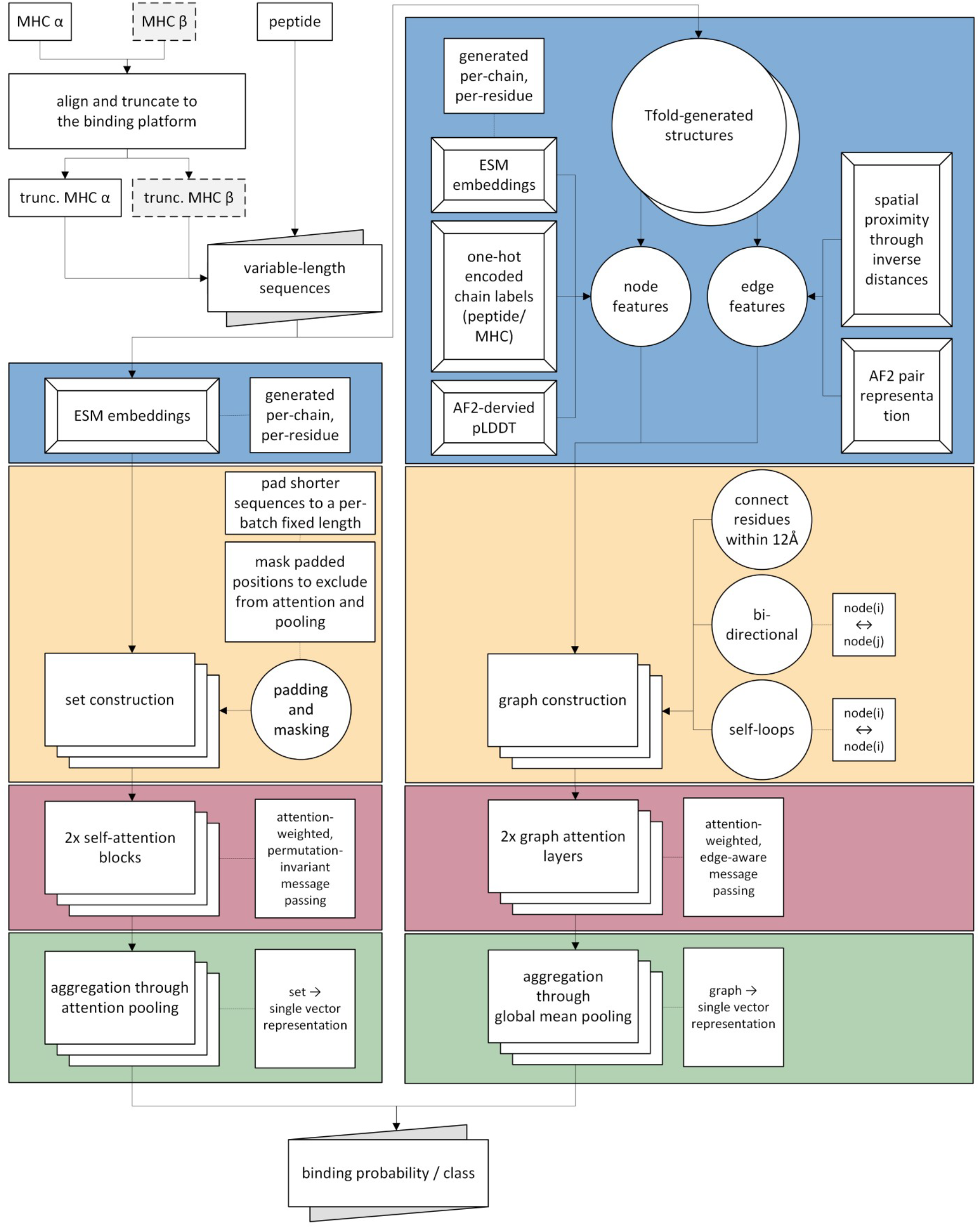
Overview of the MHChron architecture. MHC *α* sequences for classes I and II, and MHC *β* sequences for class II, are aligned and truncated to the binding platform to harmonise inputs that may or may not contain immunoglobulin-like domains. Per-residue ESM embeddings are generated for each MHC chain and peptide chain separately, then concatenated. This representation alone is used by the sequence-based model (left). In the structure-based model (right), TFold-generated structures are converted into residue-level graphs based on spatial proximity ( ≤12 Å). Per-residue node features consist of any combination of ESM embeddings, chain identity, and pLDDT scores, while pairwise edge features incorporate inverse inter-residue distances and/or AF2 pair representations. For the sequence-only framework, shorter sequences are padded to a per-batch fixed length and masked to exclude padded positions from attention and pooling. Two self-attention blocks perform attention-weighted message passing, followed by attention pooling to produce a single set-level representation that captures residue-level interactions. In the structure-based framework, graph construction is followed by two graph attention layers with edge-aware message passing, and global mean pooling to obtain a graph-level representation. In both variants, pooling yields a single fixed-dimensional representation that is passed to a final prediction layer to produce a binding probability or class label.

### 2.3 Stratified split

The stratified split enables assessment of in-distribution performance across MHC classes, alleles, and peptide lengths. On this test split, MHChron-*edgy* slightly outperformed MHChron-*light* across all metrics (AUROC 88% vs. 87%, AUPRC 89% vs. 88%, F1 score 83% vs. 82%) (Fig. 4A). Precision–recall curves were similar for both models, maintaining *>*80% precision up to ∼80% recall and substantially exceeding the baseline (54.6%) across most recall levels (Fig. 4B). Both models were reasonably well calibrated, but MHChron-*edgy* showed a closer alignment with the ideal calibration curve and a lower Brier score (0.1384 vs. 0.1465), suggesting more reliable probability estimates (Fig. 4C). Performance remained high across all peptide lengths, including atypical cases (Fig. 4D). Across loci, AUROC values ranged from 82% to 89%, with the GNN consistently matching or slightly outperforming the graph-free MHChron-*light*. Absolute AUPRC values varied between 66%–91%, reflecting differences in positive-class prevalence between loci. However, when AUPRC was normalised to the random-classifier baseline (i.e. the positive-label prevalence), both models showed strong gains, including ∼40%pt improvements for loci with lower positive-label prevalence (e.g. locus A) and near-saturation for high-prevalence loci such as DQ and DR (Fig. 4E).

**Fig. 4.**
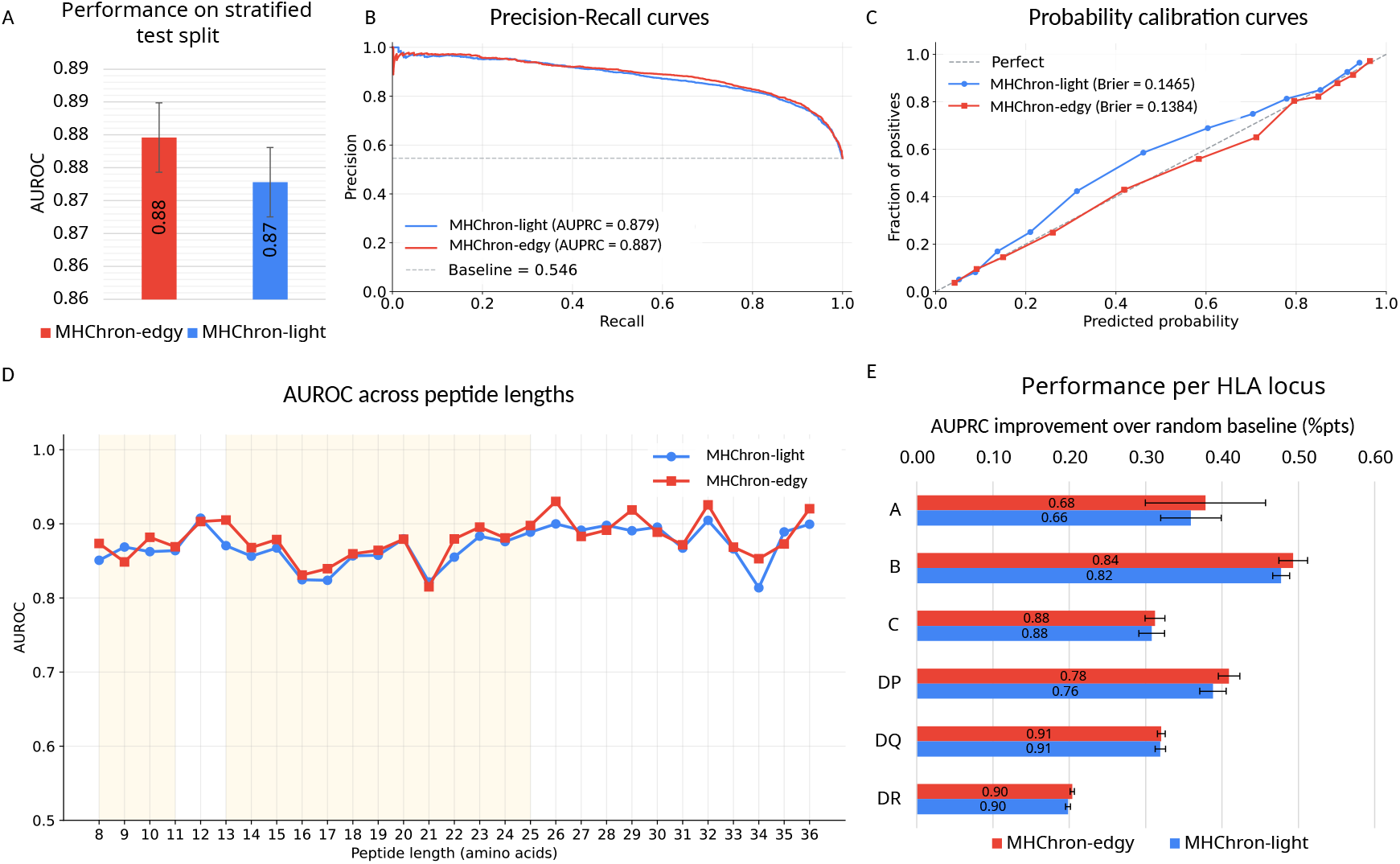
Performance on the stratified test split. A) The structure-based MHChron-*edgy* shows slightly higher AUROC than the sequence-only MHChron-*light* ; overlapping error bars indicate 95% confidence interval (CI) across five-fold cross-validation. B) Precision–recall curves show similar behaviour for both models, maintaining *>*80% precision (accuracy of positive predictions) up to approximately 80% recall (completeness of positive results) and substantially exceeding the positive-label prevalence baseline (54.6%). C) Both models are reasonably well calibrated, with MHChron-*edgy* showing closer agreement with the ideal calibration curve and a lower Brier score. MHChron-*edgy* is slightly overconfident at 0.7 predicted probability, while MHChron-*light* remains overly cautious in the range 0.1-0.8. D) Performance remains consistently high across peptide lengths, including atypical lengths (region shaded in yellow indicates the commonly observed ranges: 8-11 for MHC-I, and 13-25 for MHC-II). E) Across loci, both models show substantial improvements in AUPRC relative to locus-specific random-classifier baselines (positive-label prevalence), with MHChron-*edgy* consistently matching or exceeding the performance of MHChron-*light*. Bar heights represent the improvement over baseline (AUPRC minus baseline, in percentage points), while bar labels indicate the corresponding absolute AUPRC values. Error bars represent 95% CI of the improvement over baseline across five-fold cross-validation.

### 2.4 Leave-one-peptide-cluster-out splits

A more typical application of pMHC models involves predicting the binding of unseen peptides to previously studied alleles. Gibbs clustering [42] identified two peptide motif classes across the dataset: a charge-polarised motif with basic C-terminal anchors, and a canonical hydrophobic anchor motif dominated by aliphatic and proline residues (see Fig. SI5 for the clustering report). These clusters showed no skew with respect to MHC class, indicating that performance was not driven by differences between class I and class II molecules.

To assess whether models trained on one binding mode could generalise to the other, we performed a leave-one-peptide-cluster-out (LOPCO) evaluation, holding out each motif class in turn. Both models retained strong predictive performance when evaluated on the unseen motif, indicating generalisation beyond the dominant training pattern. MHChron-*edgy* showed a small but consistent advantage over MHChron-*light*, exceeding the sequence-only model by 1–2%pts in the charge-polarised motif (AUROC 86% vs. 85%, AUPRC 86% vs. 84%, F1 score 80% vs. 78%) and by approximately 1%pt across all metrics in the hydrophobic motif (AUROC 86% vs. 85%, AUPRC 85% vs. 84%, F1 score 79% vs. 78%).

Similar performance across motif-held-out evaluation suggests that the models learn transferable binding features rather than memorising a single peptide motif. The consistent, though modest, advantage of MHChron-*edgy* indicates that incorporating structural information improves extrapolation across divergent peptide motifs.

### 2.5 Leave-one-allele-cluster-out splits

To evaluate model generalisation to unseen MHCs while allowing peptides to have been seen during training, we performed a per-locus leave-one-allele-cluster-out (LOACO) evaluation. Across class I loci, both models retained predictive signal on unseen allele clusters, indicating generalisation beyond the allelic sequence clusters observed during training. MHChron-*edgy* consistently outperformed the sequence-only MHChron-*light*. For HLA-A, AUROC was similar for both models ( ≈64%); MHChron-*edgy* showed a larger AUPRC improvement over random baseline (+10% vs. +7%pts). The performance gap widened for HLA-B and HLA-C, where MHChron-*edgy* reached AUROCs of 72% and 77%, respectively, compared with 65% and 67% MHChron-*light*, alongside larger AUPRC gains over baseline, particularly for HLA-C (+21%pts vs. +12%pts). Averaged across all class I loci under the LOACO evaluation, MHChron-*edgy* achieved a mean AUROC of 71% and an average AUPRC improvement over random baseline of 17%pts, compared with 65% and 11% for MHChron-*light*.

In contrast, neither model generalised effectively to unseen class II allele clusters, with performance often approaching that of a random classifier. While both models exhibited modest signal for the HLA-DR locus (AUROC ≈61%, AUPRC improvement over random baseline +11%pts, F1 score 67–70%), overall predictive power remained limited, highlighting the difficulty of extrapolating to unseen MHC-II alleles.

### 2.6 Comparison with state-of-the-art (SOTA) methods

Next, we benchmarked our models against widely used external pMHC prediction tools spanning sequence- and structure-based approaches for both MHC-I and MHC-II. Specifically, we evaluated NetMHCpan-4.2 (sequence-based, class I) [75], NetMHC-IIpan-4.3 (sequence-based, class II) [33], MHCfold (structure-based, class I, 8-10mers only) [49] and MHC-II3D (structure-based, class II, HLA-DR only) [54]. External methods were evaluated as distributed and were neither retrained nor assessed for potential overlap between their original training data and the evaluation set.

This comparison was conducted using the test set of the stratified split. To ensure fair comparisons, each method was evaluated only on the subset of the test data compatible with its native input requirements. To avoid artificially inflating performance metrics, MHChron models were evaluated on exactly the same subset of samples used for each corresponding external tool. Fig. 5 reports performance in terms of AUPRC improvement over random baseline for each external tool and MHChron.

**Fig. 5.**
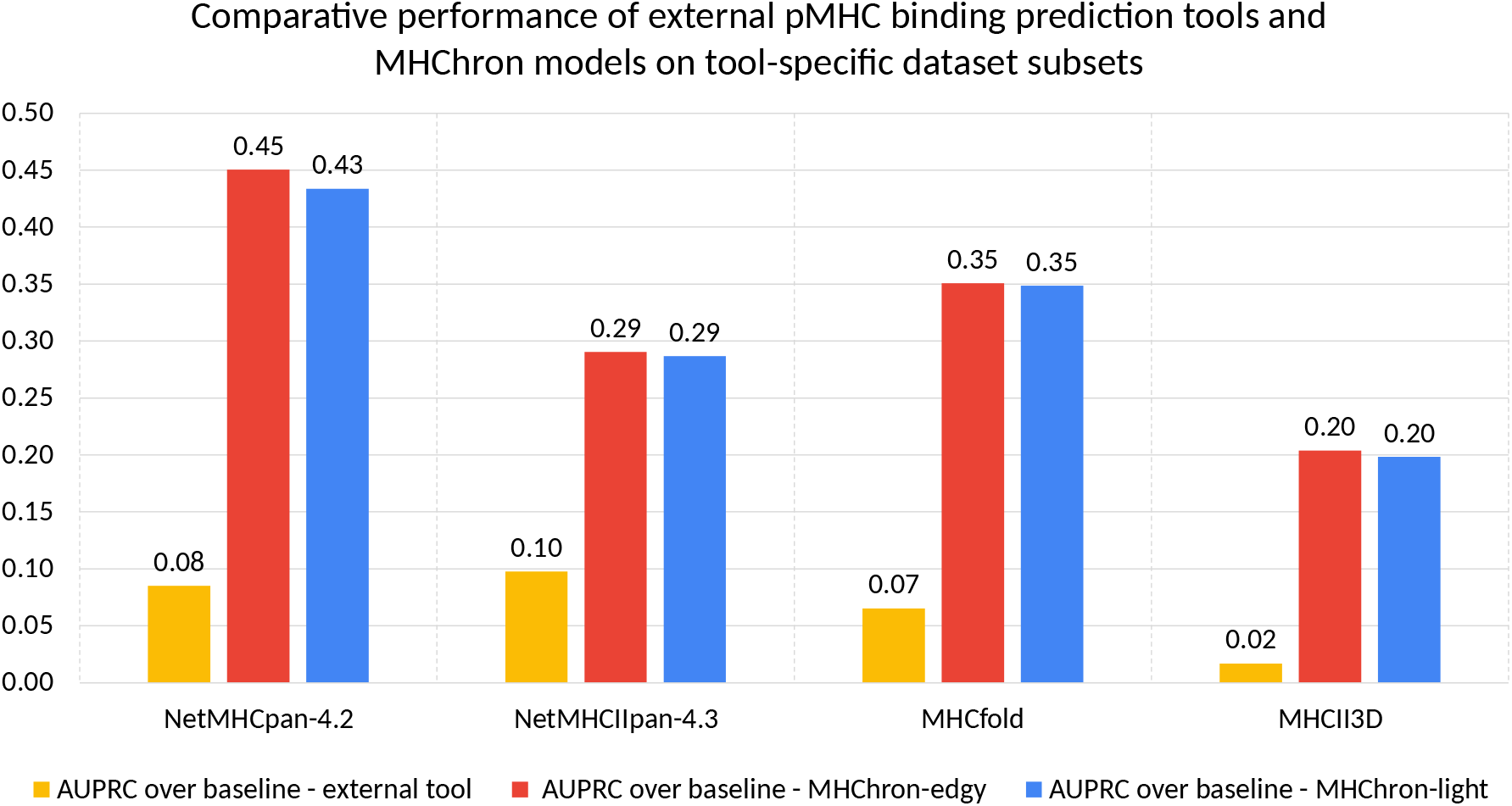
Performance comparison between MHChron and external pMHC prediction tools. External tools (yellow) include sequence-based methods (NetMHCpan-4.2, NetMHC-IIpan-4.3) and structure-based methods (MHC-fold, MHC-II3D). MHChron models include sequence-based MHChron-*light* (blue) and structure-based MHChron-*edgy* (red). Performance is compared in terms of AUPRC relative to the baseline positive rate to account for class imbalance and highlight the actual predictive gain in distinguishing binding and non-binding pMHC pairs. MHChron models consistently outperform all external baselines by a large margin.

Among external baselines, NetMHCpan-4.2 achieved an AUROC of 61% and an AUPRC of 46% (+8%pts over baseline), while NetMHC-IIpan-4.3 reached an AUROC of 64% and an AUPRC of 70% (+10%pts); MHCfold attained an AUROC of 61% and an AUPRC of 61% (+7%pts), whereas MHC-II3D achieved an AUROC of 53% and an AUPRC of 71% (+2%pts).

In contrast, both sequence-based and structure-based MHChron models outperformed all external baselines tested. The graph-based MHChron-*edgy* achieved an average AUROC of 86% and AUPRC of 88% (+32%pts average AUPRC improvement over random baseline). The graph-free MHChron-*light* also performed strongly, reaching an AUROC of 85% and AUPRC of 87% (+32%pts average AUPRC improvement over random baseline).

### 2.7 Generalisation to temporally held-out experimental data, including single-point mutants

To assess generalisation beyond the training distribution, both MHChron models were trained using all available data except for a randomly selected 7% of the stratified test split, which was held out as validation subset to stabilise training. The resulting models were evaluated on 1000 temporally held-out pMHC binding measurements deposited in the IEDB database between 16 July 2025 and 17 December 2025, which were unavailable during model development. Full and subsampled profiles of this hold-out set are presented in Fig. SI5 and SI6.

On this temporally held-out dataset, MHChron-*edgy* achieved an AUROC of 78% and an AUPRC of 94%, representing a 7%pt improvement over random baseline. MHChron-*light* achieved an AUROC of 81% and an AUPRC of 93%, representing a 6%pt improvement over random baseline. These results indicate that both models maintain discriminative performance when evaluated on newly released experimental data.

This dataset was particularly challenging because it was enriched for near-identical variants, including 181 pairs of data points (362 pMHC examples) in which the MHC sequence was identical, but peptides differed by a single amino acid, and one peptide was experimentally classified as a binder and the other as a non-binder. For each pair, a prediction was considered correct if the model assigned a higher binding probability to the experimentally binding peptide than to the non-binding peptide. Under this pairwise ranking criterion, MHChron-*edgy* and MHChron-*light* achieved accuracies of 67% and 66%, respectively, demonstrating limited but measurable sensitivity to single-residue substitutions.

## 3 Discussion

### 3.1 Balanced datasets support generalisable pMHC prediction

With some notable exceptions [41, 76–79], tools trained and evaluated on both MHC classes remain rare in pMHC binding prediction. This reflects fundamental differences between class I and class II molecules, including the number and organisation of protein chains forming the platform domain, their binding mechanisms, and the antigen-processing pathways responsible for generating their ligand repertoire. In this work, both classes were included to increase data diversity and encourage class-agnostic generalisation. To further support this objective, the combined MHC-I/II dataset was carefully engineered to ensure balanced representation across peptide lengths, allele coverage, and binding labels, while keeping it pragmatically small to facilitate extensive model development and benchmarking. Together with the rigorous data splits employed, these design choices promote the development of less biased models and better reflect real-world applications, where redundancy is limited and robust generalisation is essential.

Our results validate this approach:

- MHChron achieved consistently high metrics (AUROC ∼88%, AUPRC ∼89%, F1 ∼83%) across stratified and LOPCO splits, indicating that it is capable of effective in-distribution predictions and extrapolation to unseen peptide motifs.
- While performance declined under the LOACO split, generalisation to unseen alleles is a common challenge to pMHC binding prediction tools, and MHChron’s performance remains competitive with other SOTA methods. For MHC-I, MHChron-*edgy* ‘s performance is comparable to that reported by [80], who achieved a 22%pts improvement in AUPRC over baseline across MHC-I overall, whereas MHChron-*edgy* achieved an average improvement of 17%pts, with improvements of 10, 20, and 21%pts for the HLA loci HLA-A, HLA-B, and HLA-C, respectively. In addition, MHChron-*edgy* outperformed, and MHChron-*light* closely matched, the generalization performance of the multimodal framework proposed by [81], who reported a 12%pts improvement in AUPRC over a random baseline under a non-sequence-similarity-based rare-allele split. In contrast, generalisation to unseen class II alleles remained limited for both MHChron models, likely reflecting the greater heterogeneity and more complex binding modes of MHC-II, a challenge similarly noted by [76]. Given the breadth of the training data, which spans 312 unique MHC alleles across MHC-I and MHC-II, we anticipate that many real-life applications will involve alleles already seen during training.
- Both models outperformed all external baselines compared to in this study, with average AUPRC improvement over random baseline of ∼32%pts for MHChron, compared to an average of ∼7%pts achieved by other methods.
- Both MHChron models maintained robust performance on temporally held-out data released after model development (AUROC 81% and AUPRC 93% for MHChron-*light* ), including modest performance in ranking binding and non-binding of near-identical peptide variants.

Overall, these results demonstrate that careful dataset construction and architectural design substantially improve performance and generalisation of sequence-based pMHC predictors, while incorporating structural information provides a modest but consistent additional benefit, particularly for extrapolation to unseen alleles.

### 3.2 Sequence-based vs structure-aware modelling

The two MHChron models use the same ESM-derived per-residue features but differ in their inductive biases. MHChron-*light* employs a Set-Transformer that aggregates contextualised ESM embeddings through global self-attention to capture long-range dependencies and dominant sequence-level binding signals without imposing explicit structural connectivity assumptions. In contrast, the GATv2Conv-based MHChron-*edgy* introduces a structure-aware inductive bias by explicitly encoding local contact neighbourhoods within residue-level graphs. Their comparable expressive capacity enabled a controlled comparison between sequence-based and structure-aware approaches to pMHC binding prediction.

Prior work has shown that ESM embeddings encode substantial implicit structural and relational information, including coarse contact patterns, secondary structure tendencies, and distance constraints [22]. Consistent with this, these embeddings alone proved sufficient for strong predictive performance in MHChron-*light* and dominated the contribution of auxiliary structural inputs in MHChron-*edgy*. Furthermore, augmenting distance-based graph edges with AF2 pair representations yielded only marginal gains, suggesting considerable redundancy between these features and the information already captured by the pretrained embeddings. Together, these findings indicate that message passing primarily refines existing signal rather than introducing fundamentally new information.

One important limitation of this setup is that ESM embeddings were generated independently for the peptide and MHC chains and only concatenated subsequently. This prevents explicit cross-chain attention during representation learning, forcing inter-chain compatibility to be inferred entirely by the downstream model. The resulting lack of binding-specific interaction context may partly explain the residual advantage of the structure-aware model.

Overall, the majority of predictive signal appeared to be recoverable from residue-level representations generated by pretrained PLMs. Empirically, MHChron-*light* matched MHChron-*edgy* within ∼2% across most evaluation settings, with the gap widening primarily in LOACO, where explicit structural inductive bias improved generalisation to unseen alleles. These findings position sequence-based architectures such as the SetTransformer as competitive and computationally efficient alternatives for many pMHC binding prediction tasks, while suggesting that graph-based approaches retain value in the most challenging out-of-distribution settings.

### 3.3 Accessibility of pMHC tools

We sought to benchmark MHChron against recently published structure-based pMHC prediction methods; however, several could not be fully evaluated due to practical limitations in their publicly available implementations (i.e. unmaintained codebases, unreliable execution, or requirement for user-supplied structural templates and sequence alignments). These experiences highlight a broader challenge in the current structure-based pMHC modelling landscape: tools that rely on fragile dependencies, undocumented preprocessing steps, or specialised computational resources are inherently difficult to apply consistently across datasets and studies.

Given the closely matched performance of graph-based and graph-free MHChron models across most evaluation regimes, and the markedly lower resource demands of the latter, we deploy the sequence-based MHChron-*light* as the primary user-facing tool, available at https://github.com/wells-wood-research/MHChron. To minimise barriers to adoption, MHChron provides streamlined installation via a reproducible conda environment, minimal input requirements, and clear runtime feedback. In addition, an automated workflow for whole-protein immunogenicity scanning is included, enabling generation of overlapping peptides of user-defined lengths, evaluation against selected MHC alleles, and aggregation into interpretable per-residue binding profiles.

## 4 Conclusion

This work presents MHChron, a unified pMHC binding prediction framework that highlights the importance of balanced, diverse datasets and rigorous evaluation for generalisable pMHC binding prediction. By training on a carefully curated dataset spanning both MHC class I and II, with broad allele coverage and peptide-length diversity, MHChron moves beyond the narrow applicability regimes that limit many existing predictors.

Across progressively challenging evaluation settings, both MHChron models delivered consistently strong performance, with AUROC and AUPRC values typically in the 85%–88% range. Performance decreased under the LOACO regime, reflecting the inherent challenge of extrapolating to unseen alleles; however, MHChron remains competitive with, and in some cases outperforms, other approaches [80, 81]. Notably, it also surpassed the evaluated SOTA predictors, even when those tools were assessed solely on the allele sets and peptide-length ranges for which they were originally designed. Moreover, the deliberate emphasis on dataset balance enabled MHChron to achieve this performance while using over two orders of magnitude less training data than previously curated datasets [33, 75].

Direct comparison of sequence- and structure-based modelling revealed a trade-off: we found that the structure-aware graph-based model generalised more robustly than sequence-based models only in the most stringent regimes, but these modest gains came at a substantial computational cost.

Together, the combination of balanced data construction, broad peptide-length compatibility, and consistent performance across challenging generalisation regimes positions MHChron as a versatile and extensible resource. By coupling competitive predictive performance with practical usability, MHChron provides a foundation for future integration of additional biological signals and supports applications ranging from epitope discovery to personalised immunotherapy and protein design.

## 5 Methods

### 5.1 Dataset collection and processing

Data were obtained from publicly available peptide-binding and eluted ligand datasets (retrieval dates are provided in parentheses for online databases that are periodically updated): IEDB [64] (16/07/2025), The HLA Ligand Atlas [82] (31/07/2025), CEDAR [83] (01/08/2025), the experimentally validated VACCIMEL neoantigen dataset reported [84], and datasets associated with the development and evaluation of the following pMHC binding prediction tools: MixMHCpred3.0 [85], MixMHC2pred [86], NetMHCIIpan-4.3 [33], and NetMHCpan-4.1/NetMHCIIpan-4.0 [87]. Only linear peptides were retained; non-peptidic, discontinuous, non-classical, contaminated, or incompletely-typed entries were removed. Peptides were normalised to single-letter codes. Alleles were parsed into *α* and *β* chains (MHC-I: *α*-chain only; MHC-II: *α*/*β* chains). Rows missing required chains were discarded except HLA-DRB1/3/4/5 *β* chains, for which the missing *α* chain was assigned as HLA-DRA*01:01. Allele names were mapped to amino-acid sequences using author-supplied maps or sequences from IPD-IMGT/HLA database [64][65]; unmatched alleles (0.08%) were removed.

Binding annotations were binarised (binder=1, non-binder=0). For NetMHC-IIpan-4.3 [33] and NetMHCpan-4.1 [87], values of 0 were treated as non-binders and *>*0 as binders. Predicted non-binders without experimental confirmation were excluded. Rows mapping one peptide to multiple alleles were expanded to unique pMHC pairs. This expansion assumes the peptide binds to every associated allele; however, in eluted ligand datasets from multi-allelic sources, the specific presenting HLA is often ambiguous (i.e. the peptide may be presented by only one or a subset of the reported alleles), potentially introducing noise into the assigned labels. Reverse-orientation binders were retained without modification. After restricting to mammals and then humans (91% of the original entries), rows were deduplicated by peptide sequence and *α*/*β*-chain sequences.

Only peptide lengths with sufficient representation to enable approximately uniform sampling across lengths in the final 50,000-sample dataset were retained (7–36 aa). Due to the minimum peptide length requirements of the structure prediction tool TFold (*>*7 for MHC-I and *>*8 for MHC-II), shorter peptides were removed.

TFold truncates each input MHC sequence to the peptide-binding domain before structure prediction. Consequently, all downstream analyses after structure generation - including ESM embedding generation, dataset splitting, and model training - were performed using these truncated sequences rather than the full-length allele sequences. Because multiple alleles share identical peptide-binding domain sequences, truncation reduced the number of unique MHC sequences used in downstream analyses relative to the number of unique allele chains. The only exception was allele clustering for the exploratory dataset split, which was performed using full-length allele sequences (as described in Section SI4).

### 5.2 Decoy generation

Decoy non-binder candidates were generated by pairing each peptide with same-class alleles for which no positive binding record was available. Sampling weights were assigned to favour alleles under-represented in the dataset, and the Efraimidis-Spirakis algorithm [88] was used to select up to three unique putative negative samples per peptide.

### 5.3 Subsampling

The combined experimental and decoy dataset was reduced to a target number of examples using peptide-length-stratified subsampling. Length-specific budgets were computed by integer division with remainder reallocation, and capped where insufficient data were available. Within each peptide length, the procedure first retained all pre-specified fixed entries and positives belonging to rare allele clusters (defined as clusters with fewer than 100 positive examples). Remaining positives were sampled using Efraimidis-Spirakis weighted sampling [88], with weights inversely proportional to the number of positive examples per cluster to favour under-represented groups.

Negatives were sampled using the same weighted framework, with weights increasing with the number of positives associated with a given cluster to maintain proportionality between positive and negative examples. Experimentally determined negatives were assigned higher sampling weights than decoys, biasing selection towards original data. Global filling to reach the target number of examples was disabled to preserve per-length class balance. After concatenation and deduplication, a final trimming step was applied when necessary, again preserving fixed entries and rare positives.

### 5.4 Clustering

#### 5.4.1 Peptide clustering for the LOPCO split

Peptide sequences were clustered using GibbsCluster 2.0 [42] to identify shared binding motifs while simultaneously optimizing sequence alignment and clustering. Clustering was performed using 1-15 clusters, a fixed motif length of 9 residues, and allowing insertions and deletions of up to two residues to accommodate variable peptide lengths. Shift moves and cluster moves were enabled. Gibbs sampling was performed using the default Monte Carlo settings. For each number of clusters, the solution with the highest average Kullback-Leibler divergence (KLD) score was selected. The final number of clusters (two) was determined based on the KLD profile and the interpretability of the resulting motif classes, balancing motif specificity against potential over-partitioning.

#### 5.4.2 Allele per-locus clustering for the LOACO split

Allele sequences were first separated by locus based on the locus identifier encoded in the allele name. Within each locus, allele sequences were clustered using MMseqs2 [89] easy-cluster workflow, which applies a cascaded clustering algorithm. Clustering was performed using a minimum sequence identity threshold of 95% (– min-seq-id 0.95) with default MMseqs2 coverage and sensitivity settings. This produced 10 clusters for locus A, 17 clusters for locus B, 6 clusters for locus C, 4 clusters for DP, 17 clusters for DQ, and 11 clusters for DR.

### 5.5 Dataset split for training, validation and testing

For each split regime, a fixed held-out test set was created and used exclusively for final evaluation. The remaining data were used for model development through five-fold cross-validation, with four folds used for training and one fold used for validation in each iteration.

Three split regimes were evaluated: (1) a stratified split, which measures in-distribution performance; (2) a leave-one-peptide-cluster-out (LOPCO) split, which evaluates generalisation to peptide motif groups not observed during training; and (3) a leave-one-allele-cluster-out (LOACO) split, which evaluates generalisation to unseen allele groups. Dataset statistics for each split regime are summarised in Table 1.

**Table 1.**
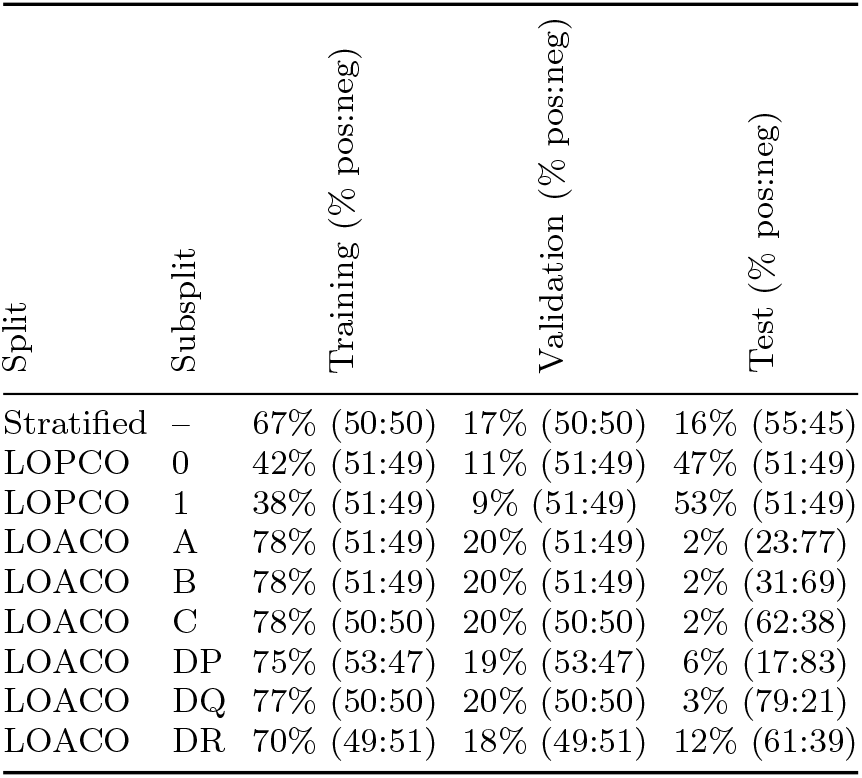
Data distribution statistics for the training, validation, and test splits, across all evaluated generalisation regimes.

Following test-set selection, the remaining data were partitioned into five cross-validation folds using a greedy cluster-aware assignment strategy. This procedure balanced fold sizes while ensuring that all examples from the same MMSeqs2 epitope cluster were assigned to the same fold, preventing closely related peptides from appearing in both training and validation sets.

#### 5.5.1 Stratified split

The stratified split was designed to estimate model performance under a representative in-distribution setting. Test examples were randomly sampled within strata defined by allele class, peptide cluster, and exact peptide length. This ensured that the test set contained examples covering different peptide lengths and allele classes rather than being dominated by the most common groups. For sufficiently large strata, the number of test examples was increased to avoid evaluating performance using very small numbers of examples. Specifically, strata containing at least 20 examples were assigned a minimum of 10 test examples where possible. The remaining examples were used for training and validation.

#### 5.5.2 LOPCO split

The LOPCO split was designed to evaluate generalisation to peptide sequence motifs not represented during training. Peptides were clustered as described in section 5.4.1, which identified two distinct motif-based peptide clusters. Evaluation was performed twice: in each experiment, one peptide motif cluster was completely withheld as the test set, while the other motif cluster was used for training and validation.

#### 5.5.3 LOACO split

The LOACO split was designed to evaluate generalisation to unseen allele groups. Alleles were clustered as described in section 5.4.2. For each locus, the largest allele cluster was selected as the held-out test set, and all examples involving alleles from this cluster were excluded from training and validation. This procedure was performed independently for each locus, resulting in separate test sets for each allele group evaluated.

### 5.6 Neural network feature generation

#### 5.6.1 Structure generation

Structures were generated with TFold [58], adjusting max pep len I/II to 36. Each pMHC was set to produce one model. Intermittent AF2 outputs, such as pair representation (d=[N res,284]) and distograms (d=[N res,N res,64]), were saved as pickle files with float32 compression.

#### 5.6.2 Per-residue and per-residue-pair feature generation

Distances were computed from AF2 distograms using dgram2dmap [90]. ESM-2 (model: esm2 t6 8M UR50D) per-residue embeddings (d=[N res,320]) were computed for peptides and each of MHC chains separately; final-layer per-residue embeddings (excluding special tokens) were concatenated and stored as float32 arrays.

#### 5.6.3 Scaler generation

For each split regime and cross-validation fold, separate StandardScaler instances (scikit-learn 1.7.2 [91]) were fitted independently for each input representation used by a model. ESM per-residue embeddings were scaled when used as node features in GNNs or as input features for the SetTransformer, and AF2 pair representations were scaled when used as edge features in GNNs. Scalers were fitted exclusively on training data from the corresponding fold. During fitting, up to 256 nodes and 3000 edges per graph were randomly sampled to reduce computational requirements. Standardisation was performed independently for each feature dimension (zero mean, unit variance), with running mean and variance estimates updated across sampled training graphs using incremental (“partial fit”) updates. The resulting scalers were then applied unchanged to the corresponding validation and test sets.

### 5.7 Neural network architecture

#### 5.7.1 GATv2Conv for graph-based MHCHron-*edgy*

To incorporate structural information, we implemented a GNN based on the GATv2Conv architecture [73] (PyTorch Geometric 2.8.0 [92]). Each pMHC complex was represented as a graph in which individual amino acids formed the nodes and edges connected all pairs of amino acids within 12^Å^. Edges were made bidirectional and self-loops were added. Node features comprised one or more of one-hot encoded peptide/MHC chain identity labels, per-position pLDDT scores, and pretrained ESM embeddings, while edge features comprised one or both of inverse pair-wise distances and AF2 pair representations. Learned input representations were standardised using precomputed StandardScaler statistics fitted exclusively on the training set (section 5.6.3).

The message-passing backbone consisted of two stacked GATv2Conv layers with 32 output channels per head, 2 attention heads, concatenated head outputs, and attention dropout of 0.3. Self-loops were disabled within the GATv2Conv layers, as they were handled explicitly during graph construction. As heads were concatenated, each layer produced 64-dimensional node embeddings. Layer normalisation and ReLU activation followed each attention layer. During training, DropEdge (p = 0.1) was applied prior to attention passes while retaining self-loops.

Node-level embeddings were aggregated into a fixed-size graph-level representation via global mean pooling. The resulting 64-dimensional graph embedding was passed to a two-layer multilayer perceptron (MLP) classifier (Linear(64 →32) →ReLU →Dropout(0.3) → Linear(32 →1)) to produce a single logit per complex.

Models were trained using binary cross-entropy (BCE) with logits and label smoothing (factor 0.05). Optimisation employed the Adam optimiser with a learning rate 1 *×* 10^−4^ and weight decay 1 *×* 10^−5^. A Reduce-on-Plateau learning-rate scheduler reduced the learning rate by a factor of 0.5 after 5 epochs without improvement in validation AUROC. Training used mixed-precision training, gradient clipping to a maximum norm of 5.0, batch size 128, and four data-loader workers. During evaluation, logits were transformed to probabilities via a sigmoid function and binarised using a threshold of 0.45.

The resulting GATv2Conv model contained 52,161 trainable parameters.

#### 5.7.2 SetTransformer for graph-free MHChron-*light*

To isolate the contribution of explicit structural information, we implemented a graph-free model based on the SetTransformer architecture [74]. The model operated directly on per-residue ESM embeddings without constructing structural edges, while matching the depth, hidden dimensionality, and training protocol of the GATv2Conv model.

Variable-length sequences were padded within each batch, and a binary mask was used to exclude padded positions from self-attention and pooling operations. Input embeddings were projected into a shared latent space using a two-layer feed-forward encoder (Linear → ReLU → Dropout(0.3) → Linear → ReLU) with a hidden dimensionality of 32. Contextualisation was performed using two stacked Self-Attention Blocks comprising multi-head self-attention (2 heads, hidden size 32) with residual connections and normalisation around both the attention sub-layer and a position-wise feed-forward network that expanded representations to four times the hidden size (128) before projection back to 32 dimensions.

A permutation-invariant attention-based pooling layer produced a 32-dimensional set representation, which was passed to a two-layer MLP classifier (Linear(32 →16) →ReLU →Dropout(0.3) →Linear(16 →1)) to generate a single logit per sequence. Training, optimisation, regularisation, and evaluation procedures were identical to those described for the GATv2Conv model.

The resulting SetTransformer model contained 41,666 trainable parameters.

### 5.8 Evaluation metrics

Predictive performance was evaluated using the area under the receiver operating characteristic curve (AUROC), the area under the precision-recall curve (AUPRC), and the harmonic mean of precision and recall (F1 score).

AUROC summarizes the trade-off between true positive and false positive rates across classification thresholds and measures the ability of the model to rank positive examples above negatives (random baseline = 0.5).

Because pMHC binding datasets are typically label-imbalanced, we additionally emphasize AUPRC, which summarizes the precision-recall trade-off and is more informative in such settings. The baseline AUPRC of a random classifier equals the prevalence of positive examples; improvements in AUPRC are therefore interpreted relative to this prevalence baseline.

Threshold-dependent performance was assessed using the F1 score. Unless otherwise stated, the classification threshold was selected on the validation set by maximizing F1 and then applied unchanged to the corresponding test set.

Performance was estimated using five-fold cross-validation. For each fold, models were trained on four partitions and evaluated on the held-out partition. Reported metrics correspond to the mean across folds, with variability reported as 95% confidence intervals based on the Student’s t-distribution.

Calibration curves were generated using 12 quantile bins to ensure approximately equal sample counts per bin.

All metrics were computed using implementations from the scikit-learn 1.7.2 library [91].

### 5.9 Testing against external baselines

#### 5.9.1 NetMHCpan-4.2 and NetMHC-IIpan-4.3

Pairs were considered strong binders if any peptide-length slice was predicted as strong; weak binders were assigned analogously. All three plausible binarisation strategies were considered for comparison: (i) treating only strong predictions as binders, (ii) treating only weak predictions as binders, and (iii) treating either strong or weak predictions as binders. The results reported in this study correspond to the first strategy (strong predictions only), which consistently yielded the best predictive performance across the evaluated metrics.

#### 5.9.2 MHCfold

Evaluation of MHCfold [49] was limited to class I test examples with peptide lengths between 8 and 10 amino acids.

#### 5.9.3 MHC-II3D

MHC-II3D [54] evaluation was restricted to HLA-DRB alleles. Allele names were normalised and converted to the tool-specific format required, and peptides were grouped by allele into per-allele input files.

Inference was performed using the official MHC-II3D binary with default parameters. The model outputs predicted IC50 values, which were used directly for evaluation. For threshold-based binary classification metrics (such as accuracy or F1-score), peptides with predicted IC50 *<* 500 nM were classified as binders. For AUROC and AUPRC computation, continuous prediction scores were defined as the negative predicted IC50 values, such that higher scores correspond to stronger predicted binding.

## Declarations

## Acknowledgements

This work was supported by a UK Research and Innovation (UKRI) funded PhD studentship through the EastBio Doctoral Training Partnership. A portion of this work was conducted during a Professional Internship for PhD Students at Aikium Inc. The authors thank Aikium Inc. for providing access to Google Cloud computing resources via cloud-computing credits, and Venkatesh Mysore and Christopher W. Wood for helpful discussions.

We acknowledge the maintainers of the Immune Epitope Database (IEDB), the HLA Ligand Atlas, and CEDAR for making publicly available immunological datasets. We thank the authors of the VACCIMEL dataset for sharing experimentally validated neoantigen data. We also acknowledge the developers of MixMHCpred, MixMHC2pred, NetMHCpan, and NetMHCIIpan for making publicly available datasets and resources that supported dataset collection and evaluation, and the developers of NetMHCpan, NetMHCIIpan, MHCfold, and MHC-II3D for making publicly available prediction tools that enabled benchmarking against state-of-the-art approaches.

## Data availability

All data supporting the findings of this study are publicly available at https://doi.org/10.5281/zenodo.18451986. The Zenodo archive contains the curated raw and subsampled pMHC datasets, prepared datasets for all cross-validation splits, datasets used for external baseline comparisons, the temporally held-out IEDB evaluation dataset, data leakage analyses confirming lack of data leakage, and the cross-validation results underlying the reported performance metrics.

## Code availability

All source code supporting this study is publicly available at .https://github.com/wells-wood-research/MHChron-paper-companion. The repository includes scripts for data collection and processing, clustering, feature generation, model training, evaluation, benchmarking against external methods, and the trained deployment models. The user-facing MHChron prediction software is available at https://github.com/wells-wood-research/MHChron.

## Author contribution

MC conceptualised and designed the study, conducted the literature review, collected and curated the data, developed and tested the neural network models, performed the formal analysis, and wrote the manuscript.

ESP substantially refactored the codebase and developed the user-facing software.

TK implemented the automated immunogenicity screening pipeline and associated visual outputs within the software.

All authors reviewed and approved the final manuscript.

