## Supplementary Information for "MHChron: diversity-balanced dataset design for robust peptide–MHC binding prediction across MHC class I and II"

### 1 Software interface

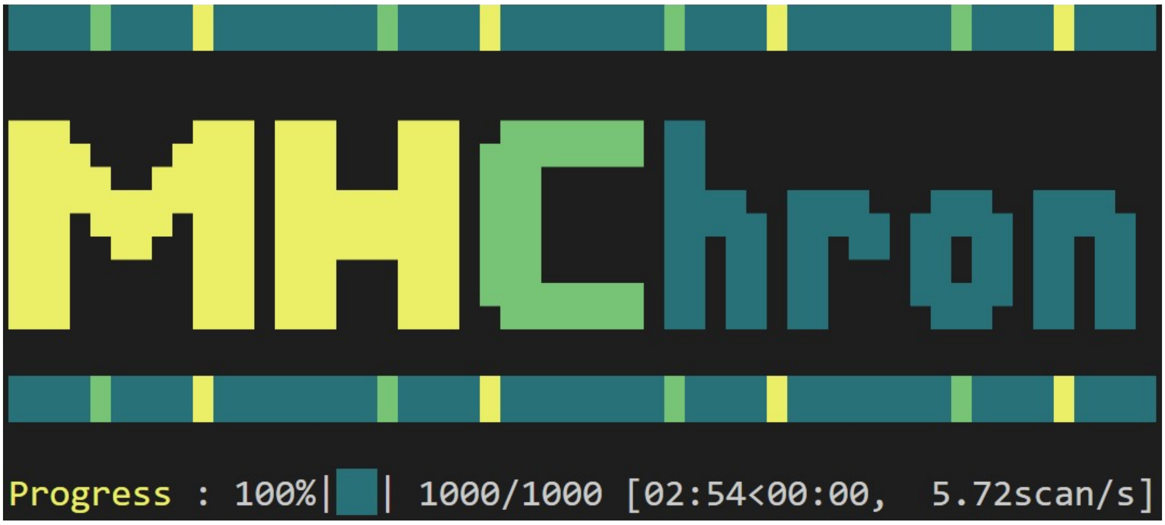


**Fig. 1 View of MHChron executed in Linux terminal**, predicting peptide-MHC (pMHC) binding on 1000 examples in less than 3 minutes.

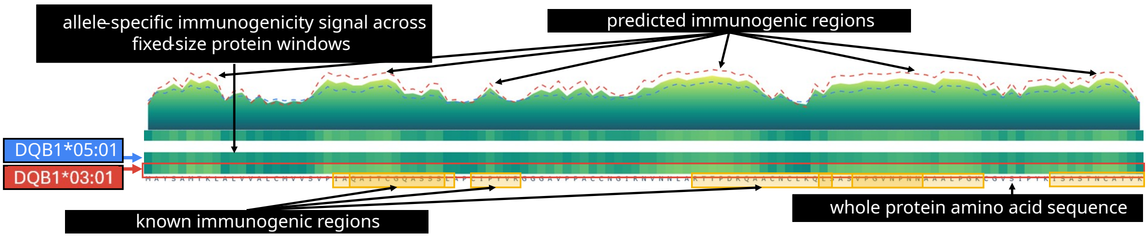


**Fig. 2** **Automated whole-protein peptide–MHC binding scanning** of the peach non-specific lipid transfer protein Pru p 3 against HLA-DQB1*03:01 and HLA-DQB1*05:01 alleles. Pru p 3 is a well-characterised food allergen with multiple experimentally validated B-cell and T-cell epitopes distributed across discrete regions of the protein sequence [1–3], providing a biologically relevant example for visualising predicted peptide–MHC binding profiles. Using fixed-length sliding peptide windows, each peptide is evaluated independently, and the predicted peptide–MHC binding probability is assigned to its corresponding sequence position. When multiple peptide lengths are analysed, predictions are aggregated to generate a continuous per-residue binding profile. In this interactive visualisation, the upper ”mountain-range” track shows the average predicted binding profile across all selected alleles, while the lower tracks display allele-specific predictions aligned to the full protein sequence. Predicted binding probability is colour-coded from yellow (high) to dark green (low). For clarity, allele-specific profiles are additionally overlaid on the averaged track as coloured dashed lines (HLA-DQB1*03:01 in red and HLA-DQB1*05:01 in blue), illustrating differences in predicted peptide binding across the same protein sequence. Regions containing experimentally reported T-cell epitopes are highlighted in yellow for qualitative comparison. Several predicted high-binding regions for HLA-DQB1*03:01 overlap or lie adjacent to these reported epitope-containing regions. HLA-DQB1*05:01 exhibits a qualitatively similar overall binding profile but generally lower predicted binding scores. This qualitative trend is consistent with published HLA association studies of Pru p 3 allergy, although such studies reflect numerous biological processes beyond peptide–MHC binding alone [4]. Importantly, none of the predicted binding peptides, either as exact matches or substrings, were present in the model training dataset, indicating that the observed agreement is unlikely to result from exact training-set memorisation.

### 2 Dataset preparation

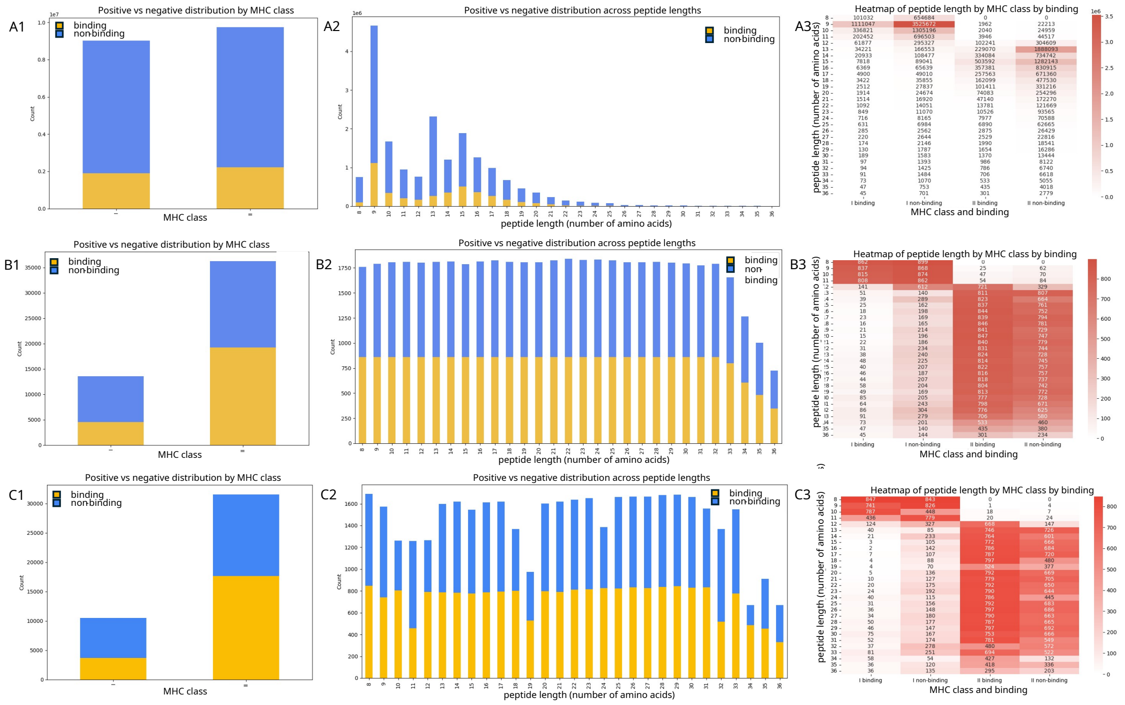


**Fig. 3 Dataset profile** of 1) MHC class representation, 2) peptide length distribution, 3) binding label distribution, across MHC class and peptide length, on A) the full dataset, including experimental and decoy information, B) the subsampled dataset, achieving perfect balance of positive and negative labels across peptide lengths and a uniform representation of all studied peptide lengths, and C) the final dataset used in subsequent model development.

### 3 Exploratory feature ablation analysis for model development

To guide model development, we performed an exploratory ablation study on a subsampled dataset using the exploratory split described in Section SI4.

Two complementary comparisons were conducted: (i) between graph-based and graph-free architectures, and (ii) between alternative node and edge feature representations within graph neural networks (GNNs). For the latter, we trained multiple GATv2Conv-based GNNs [5], each using a different controlled combination of node and edge features to quantify their contributions to predictive performance. For the former, we implemented a graph-free SetTransformer [6] with representational capacity matched to the GNN to isolate the contribution of explicit graph structure beyond the rich protein language model (PLM) embeddings (ESM) [7]. The architectures of both models are shown in Fig. 3 of the main text.

We first evaluated models using ESM per-residue embeddings as node features combined with distance-based edge features (ESM + dist), which served as a reference configuration for subsequent ablations. Building on this model, we tested the addition of per-residue pLDDT confidence scores (ESM + pLDDT + dist) and one-hot chain identity indicators (ESM + chain + dist), as well as their joint inclusion (ESM + chain + pLDDT + dist), to determine whether structural confidence and chain identity provide information beyond sequence-derived embeddings. To assess the role of alternative edge representations while keeping the node features fixed, we compared distance-based edges (ESM + dist) to AF2 pair representations alone (ESM + AF) and in combination with distances (ESM + AF + dist). Finally, to isolate the predictive capacity of non-PLM node features, we trained additional models using pLDDT-only or chain-only node features paired with either distance-based edges or AF2 pair representations.

Results for all models are summarized in terms of AUROC in Fig. SI4.

GNN models incorporating ESM per-residue embeddings consistently achieved the highest performance across all metrics (AUROC $\sim$88%, AUPRC $\sim$87%, F1 $\sim$81%), substantially outperforming models that relied solely on non-PLM node features (AUROC $<$59%, AUPRC $<$58%, F1 $<$68%). The reference ESM + distance configuration achieved an AUROC of 88.07% and an AUPRC of 86.78%, corresponding to a mean improvement of 38 percentage points over the AUPRC baseline of 49.26% determined by the positive class prevalence.

Augmenting ESM embeddings with pLDDT confidence scores or chain identity indicators yielded no improvements to performance, and no additive benefit was observed upon joint inclusion of both auxiliary features (ESM + chain + pLDDT + dist). Similarly, replacing distance-based edges with AF2 pair representations led to little improvement in performance. The ESM + AF configuration achieved an AUROC of 88.56% and an AUPRC of 87.25%, representing a marginal improvement of

0.5 percentage points over the ESM + dist model. Combining AF2 pair representations with distances (ESM + AF + dist) provided no further improvement.

The sequence-only SetTransformer model achieved an AUROC of 86.18% and an AUPRC of 84.90%, substantially outperforming models based solely on non-PLM structural features but underperforming all ESM-based GNN configurations.

Balancing predictive performance against model size, computational complexity, and feature-generation cost, we focused subsequent analyses on the ESM + distance GNN configuration and the graph-free SetTransformer model.

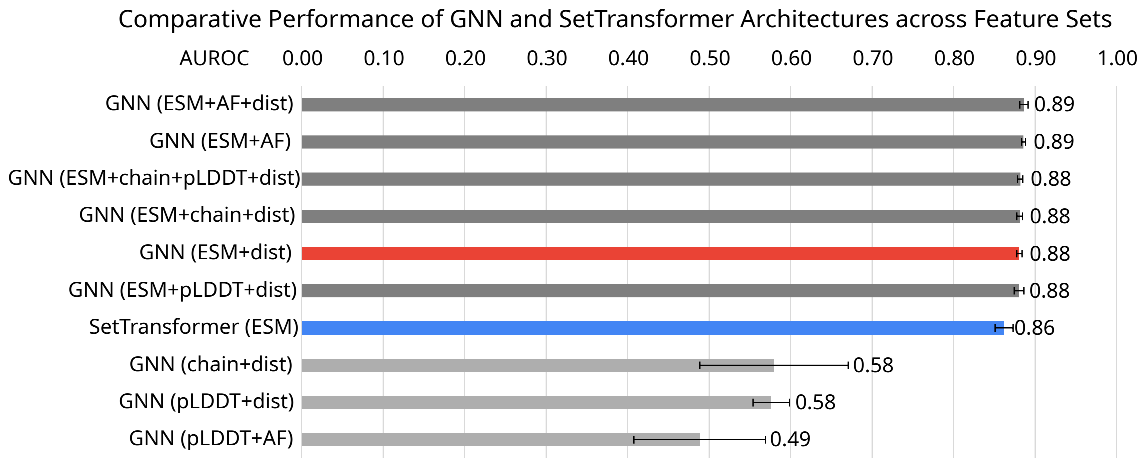


**Fig. 4 Performance comparison across model architectures and feature sets in the exploratory feature ablation analysis.** Models using ESM per-residue embeddings (top, dark grey) achieve peak performance (AUROC ≈88%), substantially outperforming models relying solely on non-PLM features (bottom, light grey). The sequence-only ESM-based SetTransformer (blue) provides a strong baseline, with the structure-embellished GNN offering modest but consistent improvements. Notably, neither the addition of pLDDT/chain identity nor using AF2 pair representations provided meaningful benefit to the reference GNN configuration (ESM + dist) (red). Results were highly stable across runs, with narrow error bars indicating 95% confidence intervals (CI) derived from the standard deviation of AUROC values across five independent cross-validation folds.

### 4 Exploratory split

In an attempt to evaluate model generalisation to unseen peptide-allele combinations, we constructed a test split by holding out entire peptide-allele cluster pairs.

Peptide and allele clusters were generated independently using MMseqs2 [8].

Allele sequences were clustered separately for the $\alpha$ and $\beta$ chains at 90% sequence identity. Clustering was performed using the full-length chain sequences, rather than sequences truncated to the platform domain; consequently, some sequences included additional immunoglobulin-like domains. This procedure produced 66 clusters from 315 unique allele chains.

Because MMseqs2 requires input sequences of at least 14 residues, peptide sequences were first lengthened by residue duplication (e.g. ABC → AABBCC). Peptides were then clustered following the protocol of [9], using an amino-acid-specific substitution matrix, high-sensitivity search, disabled low-complexity masking, a permissive E-value threshold, and increased gap penalties to discourage spurious alignments. Clustering was performed with a minimum sequence identity of 70% and the short-sequence coverage mode. To improve discrimination between short peptides, we additionally applied a custom spaced k-mer pattern (1111000011), which reduces positional correlation between k-mers and, after sequence padding, places greater emphasis on the N-terminal, central, and C-terminal positions. This procedure yielded 1,857 peptide clusters from 35,844 unique peptides.

The exploratory test split was generated by greedily selecting peptide-allele cluster pairs until the desired test-set size was reached. All observations belonging to a selected peptide-allele cluster pair were assigned to the test set, ensuring that no peptide-cluster/allele-cluster combination present in the test set appeared in the training or validation sets.

Given the high number of clusters relative to the number of unique sequences, clustering of both alleles and peptides was highly granular, with many clusters containing few sequences and limited capacity to capture broader sequence-level relatedness. As a result, this split provided only a limited challenge beyond a random partition and was therefore treated as a discovery split. It was used for exploratory model development performed before more robust splits were developed. These analyses could not be retrospectively repeated.

### 5 Peptide clustering results for the LOPCO split

### GibbsCluster report

**Version:** 2.0
**Run ID:** 973404
**Run name:** gibbs_973404
**Platform:** Linux x86_64

Read **35840 unique sequences** from file

**Settings:**
**Shift moves and cluster moves activated**
Number of clusters: 1 - 15
Motif length: 9
Initial MC temperture: 1.5
Number of temperature steps: 20
Number of iterations x Sequence x Tstep: 10
Max insertion length: 2
Max deletion length: 2
Interval between Indel moves: 10
Interval between Single Peptide moves: 20
Interval between Phase Shift moves: 100
Number of initial seeds: 1
Penalty lambda: 0.8
Weight on small clusters: 5
Preference for hydrophobic P1: 0
Sequence weighting type: 0
Background model: Uniprot pre-calculated
Use trash cluster to remove outliers: 0

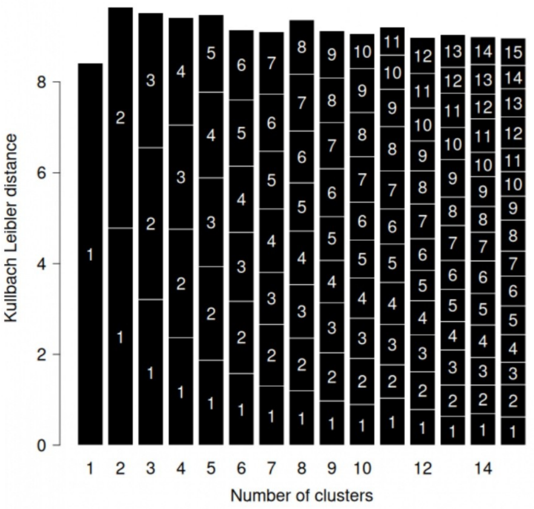


View the barplot in full size

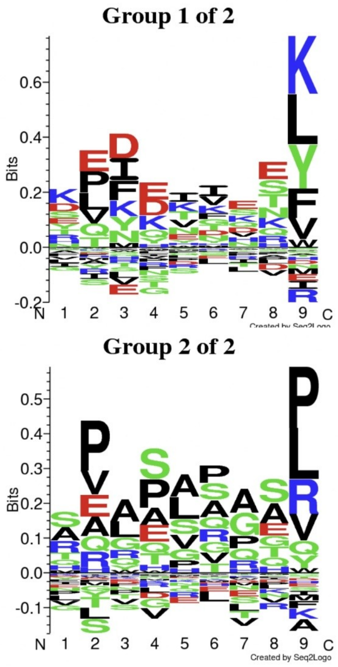


#### RESULTS for 2 CLUSTERS

Final Average KLD: **9.657746**

| Group | Size | KLD | Seq2Logo | Matrix |
|---|---|---|---|---|
| 1 | 17741 | 8.788 | Group_1of2 | Mat_1.2 |
| 2 | 18099 | 10.510 | Group_2of2 | Mat_2.2 |

Raw Clustering Report
Formatted Clustering Solution
Clustered Alignment Cores

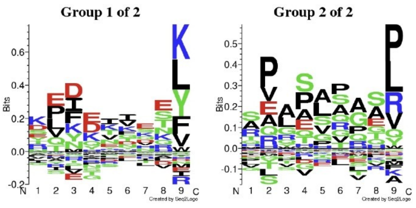


**Fig. 5 Report describing clustering of peptide sequences with GibbsCluster v2.0** [10].) Peptide sequences were clustered using GibbsCluster v2.0 [10] to identify shared binding motifs while accounting for alignment uncertainty. Clustering was performed with 1–15 clusters, a fixed motif length of 9 residues, and allowance for insertions and deletions (maximum length 2) to accommodate variable peptide lengths. Shift moves and cluster moves were enabled, and Monte Carlo sampling was run using default convergence settings. For each cluster number, the solution with the highest average Kullback–Leibler divergence (KLD) was selected. The final number of clusters (two) was chosen by comparing average KLD values across configurations, to balance motif specificity against over-partitioning. The two identified peptide motif classes represent a charge-polarised motif with basic C-terminal anchors, and a canonical hydrophobic anchor motif dominated by aliphatic and proline residues.

### 6 Dataset profile of the temporally held-out IEDB dataset

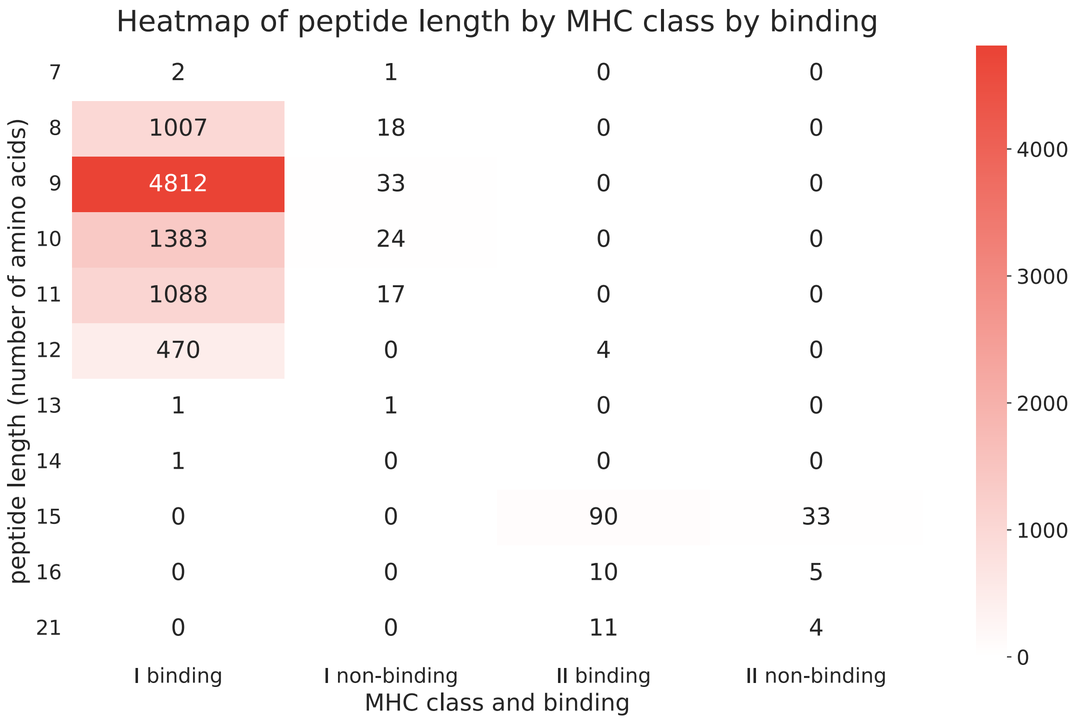


**Fig. 6 Full dataset of new experimental pMHC binding measurements** released after dataset collection for MHChron model training (IEDB database between 16 July 2025 and 17 December 2025), from which a subset of 1000 was subsampled from for testing on temporally held-out data.

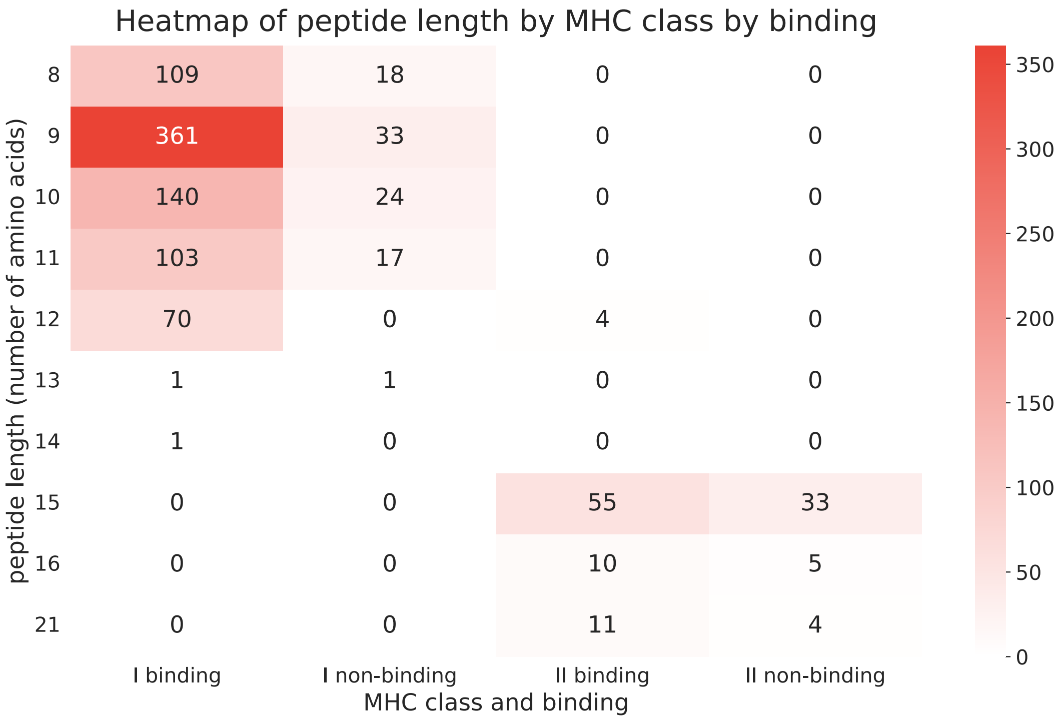


**Fig. 7 Subsampled dataset of 1000 unseen experimental pMHC binding measurements** released after dataset collection for prior model training (IEDB database between 16 July 2025 and 17 December 2025), that MHChron models were tested on.
